# Controlling Integration and Segregation in Echo State Networks via Noradrenaline and Acetylcholine Neuromodulation

**DOI:** 10.64898/2026.03.10.710755

**Authors:** Sou Nobukawa, Aya Shirama, Yusuke Sakemi, Eiji Watanabe, Teijiro Isokawa, Haruhiko Nishimura, Kazuyuki Aihara

## Abstract

Biological brain networks flexibly reconfigure functional connectivity between integration and segregation through neuromodulatory systems—noradrenaline (NA) and acetylcholine (ACh)—without altering structural connectivity. Inspired by this mechanism, we propose a modular echo state network (ESN) with context-dependent NA and ACh gain modulation, where NA promotes inter-module integration via response gain and ACh promotes intra-module segregation via multiplicative gain. We evaluate the model on two context-dependent tasks: a segregation/integration task and a context-dependent decision task. Across both tasks, the modulated model consistently outperformed the baseline, with task-appropriate modulation profiles emerging naturally from optimization and functional connectivity analysis confirming context-appropriate dynamic reorganization. These results demonstrate that neuromodulatory gain control enables adaptive, context-sensitive computation in structurally fixed reservoir networks.

## 1 Introduction

Reservoir computing (RC) is a computational framework in which input signals are projected into a recurrent neural network whose internal connections typically remain untrained, and task-relevant outputs are obtained through linear readouts from the resulting high-dimensional dynamical states [7, 16]. Owing to its ability to achieve strong performance while maintaining computational efficiency, RC has been extensively studied as an effective alternative to conventional fully trained recurrent neural networks. Among its variants, the echo state network (ESN), based on firing-rate neuron models, is one of the most representative implementations [7]. Its performance has been substantially enhanced through architectural developments such as deep ESNs [5, 6], which stack multiple sub-reservoirs to induce hierarchical dynamical responses, and assembly ESNs [19], which arrange sub-reservoirs in parallel to process multiscale high-dimensional inputs. Despite these advances, the internal connectivity of the reservoir generally remains structurally fixed, often necessitating task-specific architectural redesign to achieve optimal performance. This stands in contrast to biological brain networks, which flexibly and rapidly reconfigure their functional states without large-scale structural modifications, even under limited computational resources [8]. This discrepancy motivates the development of brain-inspired reservoir computing frameworks capable of adaptive functional modulation [3].

Biological brain networks achieve flexible and rapid reconfiguration of functional states through neuromodulatory systems, including noradrenaline (NA), acetylcholine (ACh), serotonin, and dopamine [3]. Among these, NA and ACh play central roles in dynamically regulating the balance between large-scale network integration and segregation [13]. Depending on cognitive demands and task difficulty, the brain adjusts the degree of functional integration and segregation via these modulatory systems without requiring structural rewiring. Shine and colleagues proposed a mathematical modeling framework describing such neuromodulatory effects, demonstrating that ACh activation increases the upper bound of tanh-like neuronal activation functions in targeted regions (multiplicative gain), thereby enhancing local processing, whereas NA activation steepens the slope near the activation threshold across widespread regions (response gain), thereby increasing global responsivity [13, 14]. Through these gain-modulation mechanisms, the brain dynamically shifts between segregated and integrated network states while preserving its underlying structural connectivity [2]. Such a neuromodulatory gain-control framework is highly compatible with reservoir computing, as it expands functional degrees of freedom without modifying the fixed structural connectivity of the network.

Context-dependent tasks, in which identical sensory inputs require different responses depending on contextual cues, provide a canonical paradigm for studying cognitive control [9, 10]. Theoretical and experimental studies have shown that the prefrontal cortex (PFC) implements such flexible behavior by using context signals to bias neural population trajectories rather than by altering anatomical connectivity. In addition, mixed selectivity in neural populations has been identified as a key representational principle enabling high-dimensional, context-sensitive computation in complex tasks [11]. Parallel advances in recurrent-network modeling, including biologically constrained recurrent neural networks (RNNs) and reservoir computing approaches, have demonstrated that context-dependent computation can emerge from dynamical modulation of fixed recurrent circuits [4, 15, 18]. Neuromodulation-based regulation of dynamic coordination among neuronal populations therefore represents a promising candidate mechanism for context-sensitive computation in structurally fixed networks.

In this context, we hypothesize that introducing NA- and ACh-inspired neuromodulatory mechanisms into ESNs enables dynamic regulation of the balance between integration and segregation among neuronal populations in a context-dependent manner. Specifically, we propose that functional reconfiguration—without altering structural connectivity—can support flexible, context-dependent target outputs. To test this hypothesis, we construct an ESN composed of mutually interacting sub-reservoirs and introduce NA-like modulatory signals that promote integration and ACh-like modulatory signals that promote segregation in response to contextual cues. We evaluate the proposed framework on two context-dependent tasks: (i) a segregation/integration task in which identical inputs must generate qualitatively different composite target signals depending on context, and (ii) a context-dependent decision task modelled after the paradigm of Mante et al. [9], which requires selective routing of sensory evidence based on the current context. We systematically analyze how NA and ACh modulation influence reservoir dynamics and task performance across both tasks.

## 2 Materials and Methods

### 2.1 Echo State Network with NA and ACh Neuromodulation

Previously, Shine et al. [13, 14] proposed a physiologically grounded neural network model composed of continuous neural oscillators at the population level with NA and ACh neuromodulatory signals. By adapting this framework to a discrete, modular recurrent architecture, we construct a modular leaky-integrator ESN composed of *M* sub-reservoirs (modules) {*C*_1_, *C*_2_, …, *C*_*M*_}, where module *C*_*k*_ contains *n*_*k*_ neurons, giving a total of 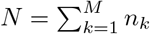 reservoir neurons. Each neuron is indexed globally by *i* ∈ {1, …, *N*}, and we write *k*(*i*) ∈ {1, …, *M*} to denote the module to which neuron *i* belongs (i.e., *i* ∈ *C*_*k*(*i*)_). In addition to the task input ***u***(*t*) ∈ ℝ^*K*^, neuromodulatory signals corresponding to NA and ACh are introduced as independent external control inputs.

#### State Update Rule

Let ***z***(*t*) ∈ ℝ^*N*^ denote the pre-activation vector:

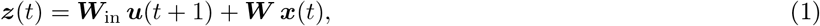

where ***W***_in_ ∈ ℝ^*N*×*K*^ is the input weight matrix. For node *i* ∈ {1, …, *N*}, the state of the NA- and ACh-modulated leaky-integrator ESN then evolves as:

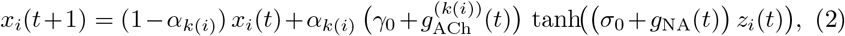

where *α*_*k*(*i*)_ ∈ (0, 1] is the leak rate of the module containing neuron *i, σ*_0_ > 0 and *γ*_0_ > 0 are baseline gain parameters, and *g*_NA_(*t*) ≥ 0 and 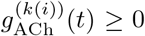 are the NA and ACh modulation levels, respectively. Assigning distinct leak rates across modules allows each module to operate at a characteristic temporal scale: a larger *α*_*k*_ yields faster dynamics suited to processing rapid sensory signals, whereas a smaller *α*_*k*_ yields slower dynamics suited to maintaining contextual information over longer intervals. The readout is computed as ***y*** (*t*) = ***W***_out_ ***x***(*t*) + ***b***, where ***W***_out_ ∈ ℝ^*L*×*N*^ and ***b*** ∈ ℝ^*L*^ are the readout weight matrix and bias trained by ridge regression, respectively.

#### Modular Reservoir Structure

The reservoir weight matrix ***W*** ∈ ℝ^*N*×*N*^ in Eq. (2) is organized into *M* × *M* blocks:

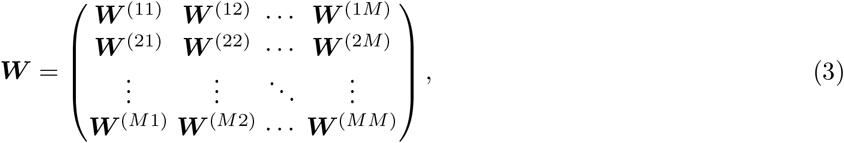

where 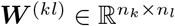 represents connections from module *C*_*l*_ to module *C*_*k*_.

Each diagonal block ***W*** ^(*kk*)^ encodes the dense intra-module connections within *C*_*k*_. A random matrix ***R***^(*k*)^ with entries drawn i.i.d. from 𝒩 (0, 1) (e.g. mean of 0 and standard deviation of 1) is generated and rescaled to a target spectral radius *ρ*_*k*_:

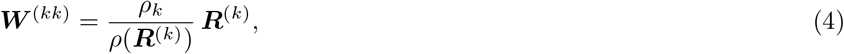

where *ρ* (***R***^(*k*)^) is the spectral radius of ***R***^(*k*)^. Setting *ρ*_*k*_ independently for each module controls the internal dynamics, including the effective time constant and memory capacity, within that module.

Each off-diagonal block ***W*** ^(*kl*)^ (*k* ≠ *l*) represents sparse inter-module connections:

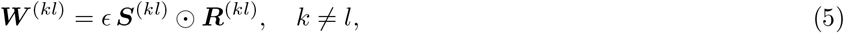

where ***R***^(*kl*)^ has entries drawn from 𝒩 (0, 1), 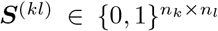 is a binary mask with each entry set independently to 1 with probability *p*_inter_ and to 0 with probability 1 − *p*_inter_, ⊙ denotes the Hadamard product, and *ϵ* > 0 controls the inter-module coupling strength. Setting *p*_inter_ ≪ 1 and *ϵ* ≪ *ρ*_*k*_ ensures that inter-module connections are sufficiently sparse and weak relative to intramodule connections.

#### Neuromodulatory Gain Control

Following Shine et al. [13, 14], NA modulates the slope of the activation function (response gain), whereas ACh scales the output amplitude of targeted populations (multiplicative gain). In Eq. (2), these effects appear as two distinct gain terms. The factor (*σ*_0_ + *g*_NA_(*t*)) multiplying *z*_*i*_(*t*) controls the slope of the tanh activation uniformly across all neurons, reflecting the broad, diffuse projections of the locus coeruleus; increasing *g*_NA_(*t*) raises the sensitivity of the entire network and promotes inter-module integration. The factor 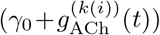 scales the output amplitude of each neuron according to its module membership, reflecting the targeted projections from the basal forebrain; increasing 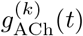 selectively amplifies the output of module *C*_*k*_ and promotes intra-module segregation.

### 2.2 Context-Dependent Multimodal Tasks

We design two context-dependent tasks to evaluate the proposed neuromodulation framework under qualitatively different computational demands. Both tasks share the three-module architecture and context-signal structure described below.

#### Common Setup

Both tasks employ a three-module ESN (*M* = 3) whose input vector ***u***(*t*) = (*u*_*s*1_(*t*), *u*_*s*2_(*t*), *c*(*t*))^T^ ∈ ℝ^3^ consists of two task-specific sensory channels and a binary context signal. To route each channel preferentially to its corresponding module, the input weight matrix ***W***_in_ ∈ ℝ^*N*×3^ is structured in block-diagonal form:

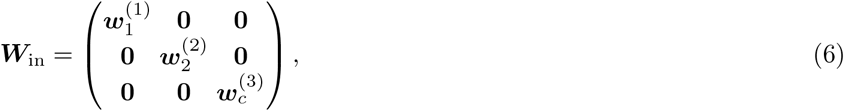

where 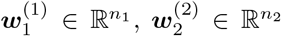, and 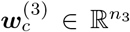 are weight vectors with entries drawn i.i.d. from 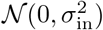, where the input scale *σ*_in_ > 0 is a tunable hyperparameter. The context signal *c*(*t*) ∈ {0, 1}, fed to the slowest module *C*_3_, is generated by partitioning the time axis into non-overlapping blocks of fixed length (context period); the context value for each block is drawn independently from a Bernoulli (1/2) distribution and held constant within the block. Feeding *c*(*t*) to *C*_3_ ensures that both the baseline and modulated models have identical access to context information, so that neuromodulation is the sole variable distinguishing the two.

#### Task 1: Segregation/Integration Task

Inspired by the multiple-timescale recurrent network framework of Yamashita & Tani [17], we design a task in which a context signal switches the required computation between local signal reproduction and cross-modal multiplication (Fig. 1(A)).

**Fig. 1.**
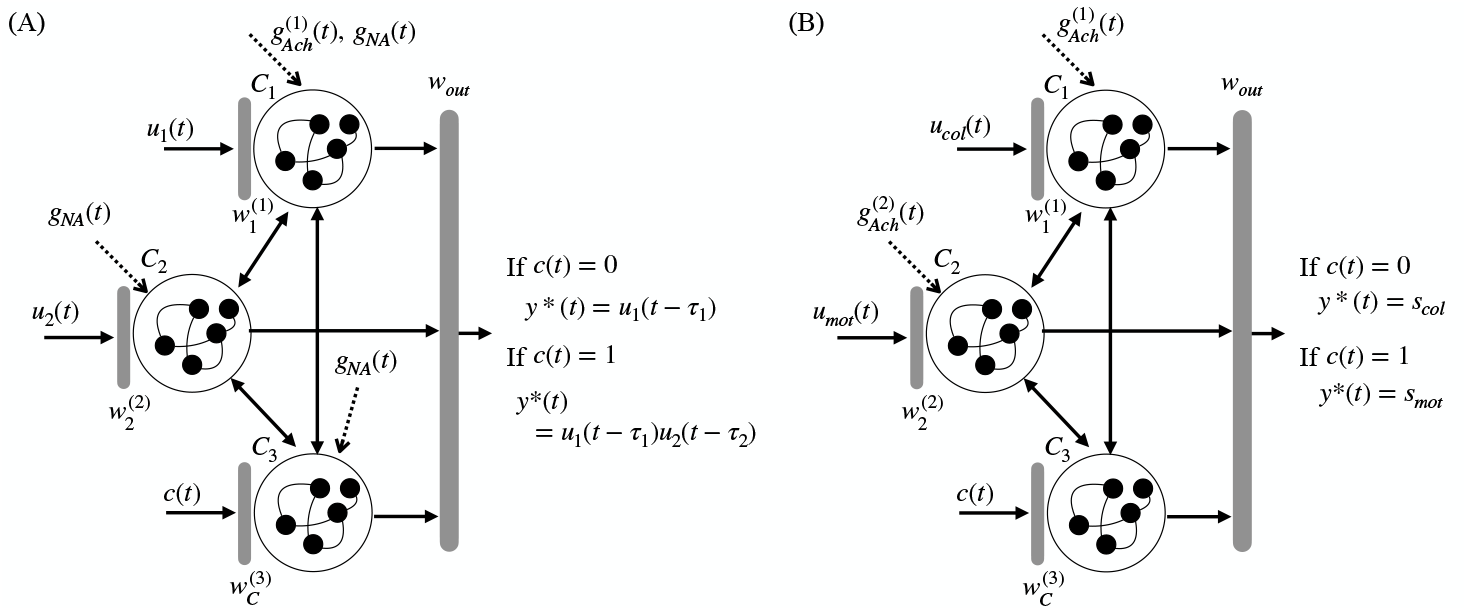
Overview of the proposed modular ESN with context-dependent NA and ACh neuromodulation. Both panels share the three-module architecture (*C*_1_, *C*_2_, *C*_3_); the context signal *c*(*t*) is fed to *C*_3_. **(A) Task 1**. *C*_1_ receives *u*_1_(*t*) and *C*_2_ receives *u*_2_(*t*). Under segregation (*c* = 0), ACh amplifies *C*_1_; under integration (*c* = 1), NA raises the response gain of all neurons. **(B) Task 2**. *C*_1_ receives *u*_col_(*t*) and *C*_2_ receives *u*_mot_(*t*). ACh selectively amplifies *C*_1_ (*c* = 0) or *C*_2_ (*c* = 1); NA is not used.

##### Input Signals

In this task the two sensory channels are set to *u*_*s*1_(*t*) = *u*_1_(*t*) and *u*_*s*2_(*t*) = *u*_2_(*t*). The first channel *u*_1_(*t*) is a fast sinusoidal signal directed to *C*_1_ (largest leak rate *α*_1_):

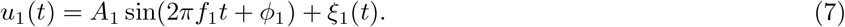

The second channel *u*_2_(*t*) is a slow sinusoidal signal (*f*_2_ ≪ *f*_1_) directed to *C*_2_:

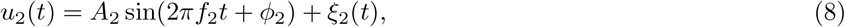

where *ξ*_1_(*t*), *ξ*_2_(*t*) ~ 𝒩 (0, 0.01^2^) are independent Gaussian observation noise terms, *A*_1_ = *A*_2_ = 0.5, *f*_1_ = 0.05 and *f*_2_ = 0.005. The context block length is set to 200 time steps (= 1*/f*_2_, one full period of *u*_2_).

##### Target Output

The target *y*\* (*t*) switches with context:

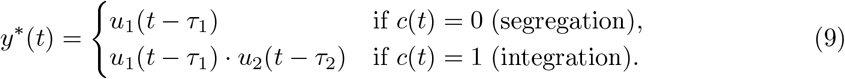

The delays were set to *τ*_1_ = 5 and *τ*_2_ = 10 time steps. Under segregation (*c* = 0), the network reproduces a delayed version of *u*_1_(*t*) using intra-module processing within *C*_1_. Under integration (*c* = 1), it must compute the product of the two delayed signals, requiring coordinated cross-modal coupling between *C*_1_ and *C*_2_.

##### Neuromodulatory Input

NA promotes inter-module integration and is activated during the integration condition; ACh promotes segregation in *C*_1_ and is activated during the segregation condition:

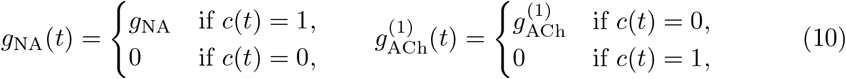

where *g*_NA_ ≥ 0 and 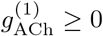 are scalar modulation levels; ACh modulation for *C*_2_ and *C*_3_ is set to zero.

#### Task 2: Context-Dependent Decision Task

This task is designed as a continuous-time analog of the context-dependent perceptual decision paradigm introduced by Mante et al. [9], in which monkeys viewed a display containing overlapping color and motion cues and had to base their binary decision on whichever attribute was designated by a context cue (Fig. 1(B)). Table 1 summarizes the correspondence between the original experimental paradigm and our computational formulation.

**Table 1.** Correspondence between the Mante et al. (2013) experiment and Task 2.

| Mante et al. (2013) | Task 2 (this work) |
| --- | --- |
| Color stream (red/green dots) | Colour input $u_{col}(t)$ directed to $C_1$ |
| Motion stream (upward/downward dots) | Motion input $u_{mot}(t)$ directed to $C_2$ |
| Stimulus coherence (S/N ratio) | Amplitude $A$ and additive noise $\xi(t)$ |
| Context cue (color or motion block) | Binary context $c(t) \in \{0, 1\}$ |
| Color context → report color | $c = 0$: target $y^*(t) = s_{col}$ |
| Motion context → report motion | $c = 1$: target $y^*(t) = s_{mot}$ |
| Selective attention in PFC | ACh gain enhancement of relevant module |

##### Input Signals

In this task the two sensory channels of the common setup are instantiated as *u*_*s*1_(*t*) = *u*_col_(*t*) (directed to *C*_1_) and *u*_*s*2_(*t*) = *u*_mot_(*t*) (directed to *C*_2_). At the start of each block of 100 time steps, two binary stimulus values *s*_col_, *s*_mot_ ∈ {−1, +1} are drawn independently and held constant for the duration of the block. The corresponding noisy sensory inputs are:

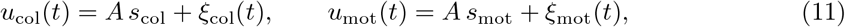

where *A* = 0.3 is the signal amplitude and *ξ*_col_(*t*), *ξ*_mot_(*t*) ~ 𝒩 (0, 0.2^2^) are independent Gaussian noise terms. The channel *u*_col_(*t*) is directed to *C*_1_ and *u*_mot_(*t*) to *C*_2_, paralleling the separate color and motion processing streams of the Mante et al. paradigm.

##### Target Output

The target output is the binary decision value of the context-relevant stream, held constant within each block:

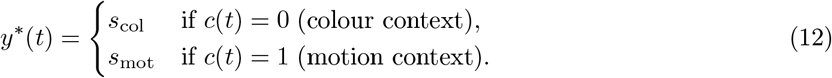

The ESN must thus suppress the irrelevant stream and selectively maintain the decision variable of the relevant stream against the corrupting noise—mirroring the selective integration of evidence observed in monkey PFC by Mante et al.

##### Neuromodulatory Input

Because the relevant stream switches with context, ACh selectively amplifies the output of the module receiving the currently relevant sensory channel; NA modulation is not used in this task (*g*_NA_(*t*) ≡ 0):

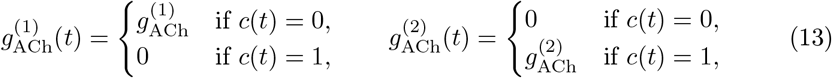

where 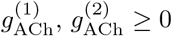, are optimized independently. This ACh-mediated selective amplification directly models the context-dependent attentional gating reported in PFC: by boosting the gain of the relevant module, the network is biased toward integrating the task-relevant sensory evidence while the competing stream is relatively suppressed.

### 2.3 Experimental Settings, Optimization Protocol, and Evaluation Metrics

The reservoir comprised *M* = 3 modules of *n*_*k*_ = 100 neurons each (*N* = 300 in total), with a washout period of 200 time steps. All remaining hyperparameters were optimized using Optuna [1] with a Tree-structured Parzen Estimator sampler over 1000 trials per condition; convergence was confirmed by inspecting the running best objective across all trials. Each trial was evaluated by averaging across 10 independent random seeds to reduce variance due to random reservoir initialization. The final performance comparison between the two models was carried out over 20 independent random seeds. The search ranges are summarized in Table 2.

**Table 2.** Fixed parameters and hyperparameter search ranges used in the numerical experiments. Common parameters are shared across both tasks; task-specific parameters apply only to the modulated model of the respective task.

| Parameter | Fixed / Search range |
| --- | --- |
| **Fixed (both tasks)** |  |
| Number of modules $M$ | 3 |
| Module size $n_k$ | 100 each ($N = 300$) |
| Washout period | 200 steps |
| **Common search range (Optuna, both tasks)** |  |
| Spectral radius $\rho$ | [0.5, 1.5] |
| Leak rate $\alpha_1$ (fast) | [0.50, 1.00] |
| Leak rate $\alpha_2$ (mid) | [0.20, 0.45] |
| Leak rate $\alpha_3$ (slow) | [0.05, 0.19] |
| Inter-module connection probability $p_{\text{inter}}$ | [0.01, 0.12] |
| Inter-module coupling strength $\epsilon$ | [0.01, 0.12] |
| Input scale $\sigma_{\text{in}}$ | [0.02, 0.25] |
| Baseline response gain $\sigma_0$ | [0.7, 1.4] |
| Baseline multiplicative gain $\gamma_0$ | [0.7, 1.4] |
| Ridge regression coefficient $\lambda$ | $[10^{-5}, 10^{-2}]$ (log scale) |
| **Task-specific search range (modulated model only)** |  |
| *Task 1: Segregation/Integration* |  |
| NA modulation $g_{\text{NA}}$ | [0.0, 1.5] |
| ACh modulation $g_{\text{ACh}}^{(1)}$ | [0.0, 1.5] |
| *Task 2: Context-Dependent Decision* |  |
| ACh modulation $g_{\text{ACh}}^{(1)}$ (colour stream) | [0.0, 1.5] |
| ACh modulation $g_{\text{ACh}}^{(2)}$ (motion stream) | [0.0, 1.5] |

To evaluate the benefit of neuromodulation fairly, a *baseline* model (with *g*_NA_(*t*) = 0 and 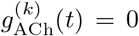 for all *t*) and a *modulated* model (with context-dependent switching as defined in Eq. (10)) were optimized independently and subsequently compared under identical evaluation conditions. Each optimization trial used a strict three-phase data split to prevent information leakage: readout weights ***W***_out_ and bias ***b*** were fitted by ridge regression on a training sequence of length *T*_train_, the objective function was evaluated on a separately generated validation sequence of length *T*_val_, and the final performance was measured on a held-out test sequence of length *T*_test_. The lengths were set to *T*_train_ = 2000 and *T*_val_ = 1200 during optimization, and *T*_train_ = 3000 and *T*_test_ = 2000 during the final comparison. The search ranges for the three leak rates were designed to be non-overlapping (Table 2), thereby guaranteeing *α*_1_ ≥ *α*_2_ ≥ *α*_3_ throughout the optimization and ensuring that *C*_1_, *C*_2_, and *C*_3_ operate at progressively slower temporal scales. The objective function differed between the two tasks. For Task 1, in which the integration condition is substantially harder than the segregation condition, the objective based on the normalized mean squared error (NMSE) was defined as: ℒ_1_ = 0.7 NMSE_all_ + 0.3 NMSE_int_, where the additional weight on NMSE_int_ compensates for its greater difficulty. For Task 2, in which the colour and motion conditions are symmetric, the objective was: ℒ_2_ = NMSE_all_, where NMSE_all_ is the NMSE averaged across both the colour and motion conditions. Trials exhibiting divergent behavior were assigned a large penalty to favor numerically stable solutions. After optimization, the best-performing parameters of each model were fixed and both models were evaluated across multiple random seeds; performance is reported as the mean and standard deviation of the condition-specific and overall NMSE values.

#### Evaluation Metrics

Task performance was quantified by NMSE computed separately for each context condition. For Task 1, this yields NMSE_seg_ (segre-gation, *c*(*t*) = 0) and NMSE_int_ (integration, *c*(*t*) = 1). For Task 2, this yields NMSE_col_ (colour context, *c*(*t*) = 0) and NMSE_mot_ (motion context, *c*(*t*) = 1). In both cases, NMSE_all_ denotes the NMSE averaged across all time steps regardless of context.

To assess the modular organization of reservoir dynamics, functional connectivity was characterized by the Pearson correlation *C*_*ij*_ between the state time series of each pair of neurons *x*_*i*_ and *x*_*j*_. Module-level connectivity was summarized by the mean absolute correlation between modules *k* and *m*:

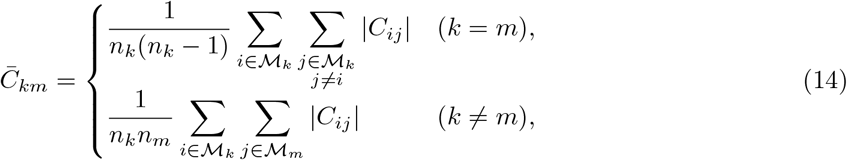

where ℳ_*k*_ denotes the index set of neurons in module *C*_*k*_ and *n*_*k*_ = |ℳ_*k*_|. The diagonal entries (*i* = *j*) are excluded from the intra-module average to avoid self-correlation.

## 3 Results

Figure 2 shows representative time series of the context signal *c*(*t*), the sensory inputs *u*_1_(*t*) and *u*_2_(*t*), and the model outputs *y* (*t*) alongside the target *y*\*(*t*) for both the baseline (no modulation) and modulated models for Task 1. The context signal *c*(*t*) switches randomly between the segregation condition (*c* = 0) and the integration condition (*c* = 1) at fixed intervals of 200 time steps. During the segregation condition, the target is a delayed version of *u*_1_(*t*); during the integration condition it becomes the product of the delayed *u*_1_(*t*) and *u*_2_(*t*), which produces a signal with substantially richer temporal structure and larger dynamic range. In both models, *y*(*t*) broadly tracks the target *y*\*(*t*) throughout, confirming that the modular ESN architecture supports context-dependent computation: the baseline relies solely on the context input *c*(*t*) fed to module *C*_3_, while the modulated model additionally exploits context-switched neuromodulatory signals and clearly outperforms the baseline in the majority of context windows, as confirmed by the per-block NMSE panel (bottom row). Figure 3 shows the corresponding results for Task 2 (Context-Dependent Decision Task). As *c*(*t*) alternates between the colour context (*c* = 0) and the motion context (*c* = 1), the target *y*\* (*t*) switches between *u*_col_(*t*) and *u*_mot_(*t*), respectively. In the baseline model, the strong additive noise leads to noticeable deviation of *y* (*t*) from *y*\*(*t*); in the modulated model, ACh-mediated selective amplification of the task-relevant input stream yields substantially higher agreement with the target across both context conditions.

**Fig. 2.**
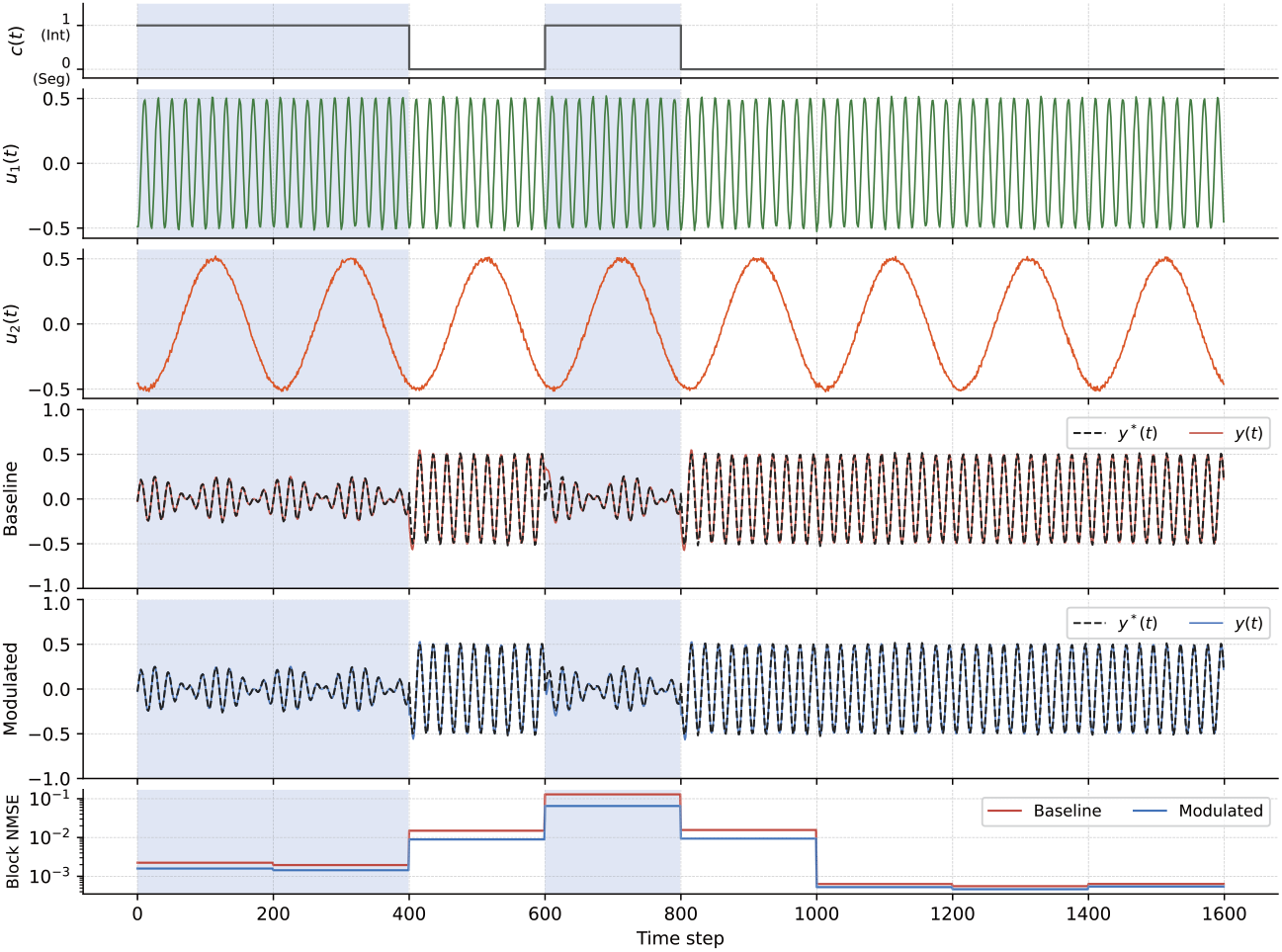
Representative time series for the baseline (no neuromodulation) and modulated (context-dependent NA and ACh) models in Task 1 (Segregation/Integration Task). Rows from top: context signal *c*(*t*); fast sensory input *u*_1_(*t*); slow sensory input *u*_2_(*t*); baseline output *y* (*t*) and target *y*\* (*t*); modulated output *y* (*t*) and target *y*\* (*t*); per-block NMSE (200-step window). Light-blue shading indicates the integration condition (*c* = 1); unshaded regions correspond to the segregation condition (*c* = 0).

**Fig. 3.**
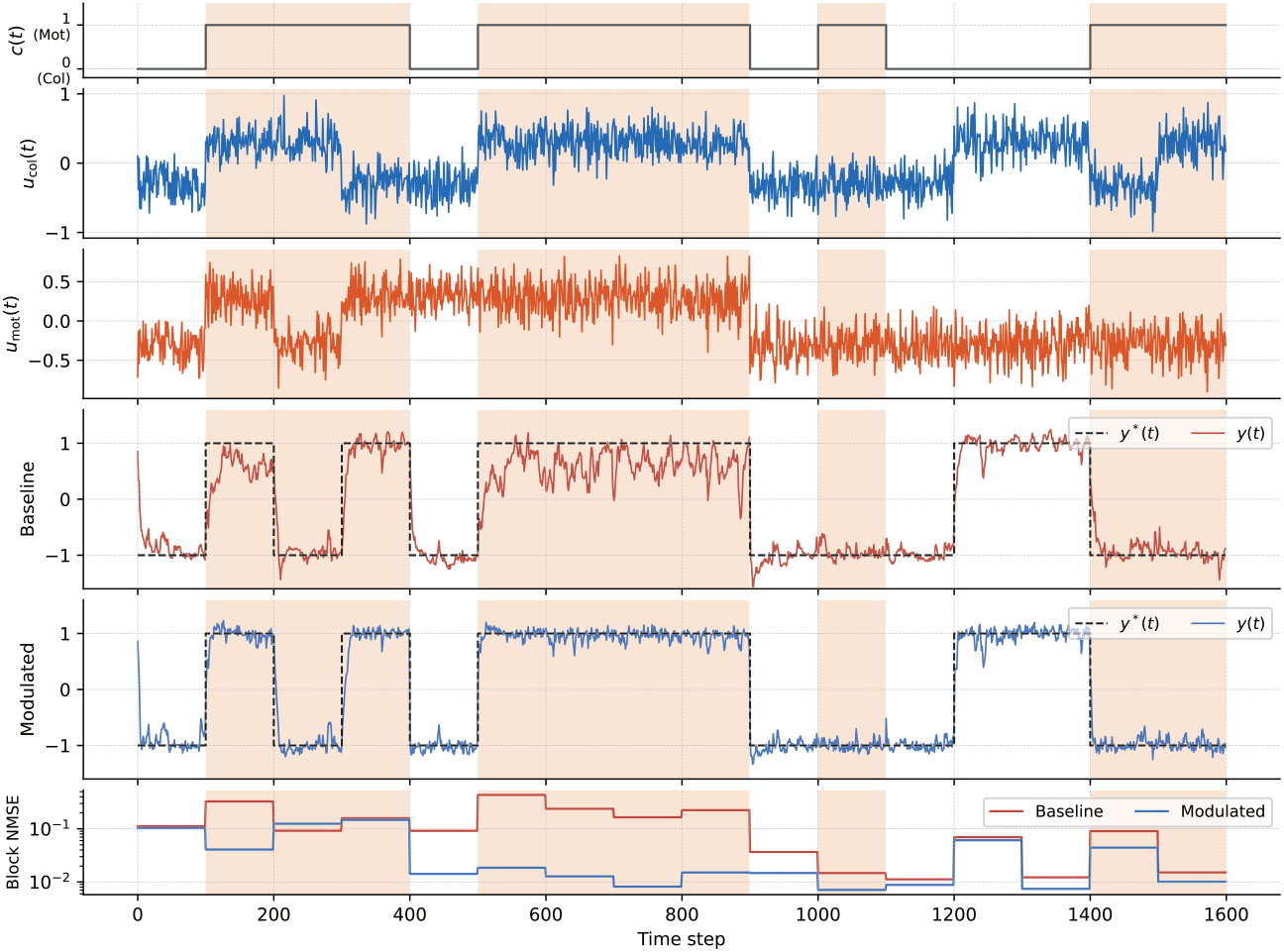
Representative time series for the baseline (no neuromodulation) and modulated (context-dependent NA and ACh) models in Task 2 (Context-Dependent Decision Task). Rows from top: context signal *c*(*t*); colour input *u*_col_(*t*); motion input *u*_mot_(*t*); baseline output *y* (*t*) and target *y*\* (*t*); modulated output *y* (*t*) and target *y*\* (*t*); per-block NMSE (100-step window). Light-orange shading indicates the motion context (*c* = 1); unshaded regions correspond to the colour context (*c* = 0).

Table 3 quantifies this observation for both tasks. For Task 1, the modulated model achieved relative improvements of 20.0% in the segregation condition (NMSE_seg_), 42.9% in the integration condition (NMSE_int_), and 30.0% overall (NMSE_all_), confirming that both ACh- and NA-mediated gain modulation are effective across context conditions. For Task 2, the improvements were similarly pronounced: 45.1% for the colour condition (NMSE_col_), 54.2% for the motion condition (NMSE_mot_), and 49.0% overall, confirming that context-dependent ACh gain control effectively suppresses irrelevant sensory streams across both context conditions.

**Table 3.** Task performance comparison (mean ± std over 20 seeds).

|  | Baseline | Modulated | Rel. imp. |
| --- | --- | --- | --- |
| **Task 1** (Segregation/Integration) |  |  |  |
| NMSE<sub>seg</sub> | 0.0147 ± 0.0136 | 0.0117 ± 0.0145 | 20.0% |
| NMSE<sub>int</sub> | 0.161 ± 0.194 | 0.0919 ± 0.110 | 42.9% |
| NMSE<sub>all</sub> | 0.0271 ± 0.0236 | 0.0189 ± 0.0194 | 30.0% |
| **Task 2** (Context-Dependent Decision) |  |  |  |
| NMSE<sub>col</sub> | 0.0863 ± 0.0448 | 0.0474 ± 0.0173 | 45.1% |
| NMSE<sub>mot</sub> | 0.0989 ± 0.0573 | 0.0453 ± 0.0135 | 54.2% |
| NMSE<sub>all</sub> | 0.0844 ± 0.0266 | 0.0431 ± 0.00860 | 49.0% |

Through optimization with Optuna, the best hyperparameters were identified for each model and task. In the Task 1 modulated model, the NA gain was *g*_NA_ = 0.719 and the ACh gain was 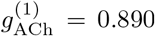 In the Task 2 modulated model, NA was inactive (*g*_NA_ ≡ 0) and ACh gains were identified: 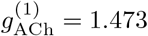 and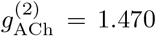, reflecting the symmetric selective-attention nature of that task. For Task 1, the optimized *ϵ* was comparable between the baseline and modulated models (0.112 vs. 0.114), indicating no structural difference in intermodule coupling. For Task 2, the modulated model adopted a substantially lower inter-module coupling strength *ϵ* compared with its baseline (0.012 vs. 0.103), suggesting that ACh gain control reduces the need for strong structural intermodule connectivity.

Table 4 shows the module-level functional connectivity 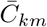 for both tasks. For Task 1, inter-module connectivity between the two sensory modules showed no large reorganization overall, but a modest condition-dependent trend was observed in the modulated model: 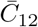 was slightly higher under the integration condition than under the segregation condition (0.401 vs. 0.351), whereas the baseline showed no such difference (0.318 vs. 0.321; Table 4). This pattern is consistent with NA-mediated response-gain enhancement facilitating cross-module coupling specifically when integration is required. For Task 2, the effect of ACh modulation was markedly different: inter-module connectivity was substantially reduced in the modulated model relative to the baseline (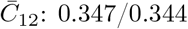 in baseline vs. 0.206/0.218 in modulated across colour/motion contexts), consistent with ACh-mediated selective amplification of the task-relevant input stream effectively isolating the relevant module from competing cross-module inputs.

**Table 4.**
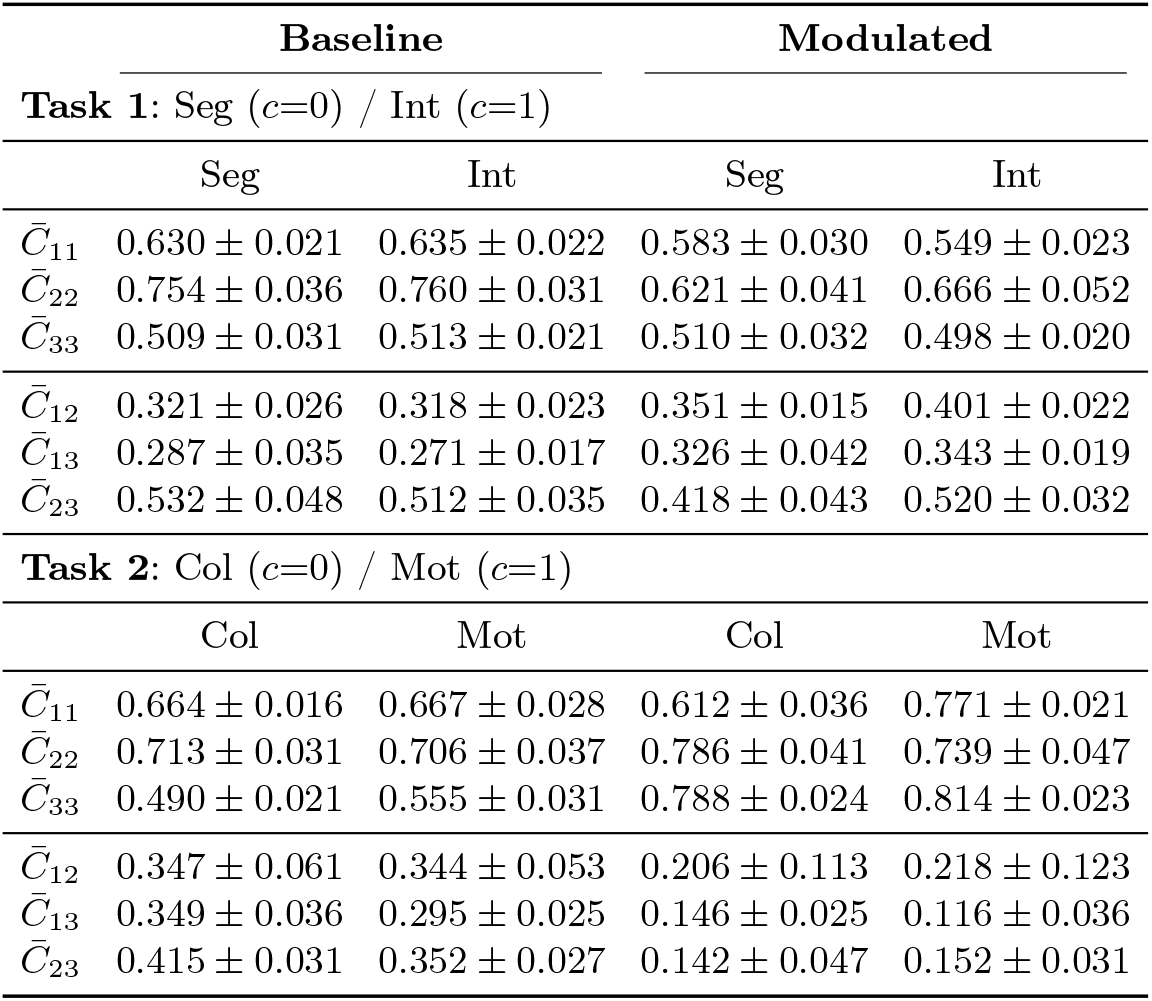
Module-level functional connectivity 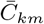 (mean ± std over 20 seeds). Diagonal: intra-module; off-diagonal: inter-module.

## 4 Discussion and Conclusions

In this study, we constructed a modular ESN incorporating NA and ACh gain-modulation mechanisms and evaluated its performance on two context-dependent tasks: a segregation/integration task (Task 1) and a context-dependent decision task (Task 2). Across both tasks, context-dependent neuromodulation consistently outperformed the unmodulated baseline; functional connectivity analysis confirmed context-appropriate reorganization of inter-module coupling in both tasks, with a more pronounced shift in Task 2. The required modulation profile differed qualitatively between tasks—Task 1 recruited both NA and ACh, whereas Task 2 relied on ACh alone?demonstrating that the neuromodulatory repertoire is naturally matched to the computational demands of each task.

We now consider the mechanistic basis for these improvements. In Task 1, ACh modulation during the segregation condition selectively amplifies the output of module *C*_1_ via multiplicative gain, strengthening the contribution of *u*_1_(*t*)-driven intra-module activity relative to cross-module inputs leaking through the sparse off-diagonal blocks of ***W***, thereby promoting functional segregation within the reservoir. During the integration condition, NA modulation increases the slope of the tanh activation function uniformly across all reservoir neurons (response gain), raising the overall network sensitivity and facilitating the reproduction of the target product output. Functional connectivity analysis for Task 1 revealed a modest but interpretable condition-dependent pattern in the modulated model: 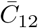 was higher under the integration condition than under the segregation condition (0.401 vs. 0.351; Table 4), whereas the baseline showed no such difference (0.318 vs. 0.321). This selective increase in inter-module coupling under integration is consistent with NA-mediated response-gain enhancement facilitating cross-module information flow when it is computationally required, and is further supported by the 42.9% improvement in NMSE_int_. The optimizer identified non-zero values for both gain parameters (*g*_NA_ = 0.719 and 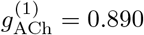): had either gain been ineffective, the optimizer would have driven it toward zero. In Task 2, ACh modulation alternately amplifies the gain of *C*_1_ (colour context, *c* = 0) or *C*_2_ (motion context, *c* = 1), selectively enhancing the representation of the task-relevant sensory stream. This mechanism is consistent with the marked reduction in inter-module connectivity in the modulated model (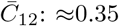 in baseline vs. ≈0.21 in modulated; Table 4), which reflects the functional isolation of the task-relevant module from cross-module interference. These gain-modulation mechanisms were previously modeled in the context of recurrent neural network frameworks by Shine et al. [14], and the present work represents their first systematic introduction within a reservoir computing framework. This approach is further motivated by recent findings from Deco et al. [3], who showed that incorporating neuromodulatory signals into whole-brain computation models is essential for capturing a broad repertoire of flexible, task-dependent behaviors—a requirement that fixed structural connectivity alone cannot satisfy.

Several directions remain for future investigation. First, the proposed framework should be evaluated on more demanding and practically relevant tasks, such as online control and multimodal sequence learning. Second, while the modulation signal is explicitly provided in this study, developing an effective method for autonomously learning to supply such signals is itself an important open problem. Although neural modulation dynamics were introduced into the reservoir in our previous study [12], that approach required substantial training cost due to reliance on backpropagation through time, and a more efficient alternative is therefore desirable. Third, additional neuromodulatory systems—including serotonin and dopamine—should be incorporated, and their interactions with NA and ACh explored, as such multi-system modulation may further enrich the repertoire of context-dependent computations achievable within the reservoir computing framework.

In conclusion, we proposed a modular ESN with context-dependent NA and ACh neuromodulation and demonstrated its effectiveness on two qualitatively different context-dependent tasks. The modulated model consistently outperformed the unmodulated baseline in output accuracy, with functional connectivity analysis confirming context-appropriate reorganization of inter-module coupling. These results establish neuromodulatory gain control as a versatile and biologically grounded mechanism for adaptive, context-sensitive computation in structurally fixed reservoir networks.

## Acknowledgment

This work was supported in part by the New Energy and Industrial Technology Development Organization (NEDO) under Project JPNP14004. This work was supported by JSPS KAKENHI, Grant-in-Aid for Scientific Research (B) (Grant Number JP25K03198, SN; 25K00148, YS) and Transformative Research Areas (A) (Grant Number JP25H02626, SN).

